# A Systematic Strategy for Identifying Causal Single Nucleotide Polymorphisms and Their Target Genes on Juvenile Arthritis Risk Haplotypes

**DOI:** 10.1101/2022.03.07.483040

**Authors:** Kaiyu Jiang, Tao Liu, Susan Kales, Ryan Tewhey, Dongkyeong Kim, Yungki Park, James N. Jarvis

## Abstract

**Introduction:** Although genome-wide association studies (GWAS) multiple regions conferring genetic risk for juvenile idiopathic arthritis (JIA), we are still faced with the task of identifying the single nucleotide polymorphisms (SNPs) on the disease haplotypes that exert the biological effects that confer risk. Until we identify the risk-driving variants, identifying the genes influenced by these variants, and therefore translating genetic information to improved clinical care, will remain an insurmountable task. We used a function-based approach for identifying causal variant candidates and the target genes on JIA risk haplotypes.

**Methods:** We used a massively parallel reporter assay (MPRA) in myeloid K562 cells to query the effects of 5,226 SNPs in non-coding regions on JIA risk haplotypes for their ability to alter gene expression when compared to the common allele. The assay relies on 180 bp oligonucleotide reporters (“oligos”) in which the allele of interest is flanked by its cognate genomic sequence. Barcodes were added randomly by PCR to each oligo to achieve >20 barcodes per oligo to provide a quantitative read-out of gene expression for each allele. Assays were performed in both unstimulated K562 cells and cells stimulated overnight with interferon gamma (IFNg). As proof of concept, we then used CRISPRi to demonstrate the feasibility of identifying the genes regulated by enhancers harboring expression-altering SNPs.

**Results:** We identified 553 expression-altering SNPs in unstimulated K562 cells and an additional 490 in cells stimulated with IFNg. We further filtered the SNPs to identify those plausibly situated within functional chromatin, using open chromatin and H3K27ac ChIPseq peaks in unstimulated cells and open chromatin plus H3K4me1 in stimulated cells. These procedures yielded 42 unique SNPs (total = 84) for each set. Using CRISPRi, we demonstrated that enhancers harboring MPRA-screened variants in the *TRAF1* and *LNPEP/ERAP2* loci regulated multiple genes, suggesting complex influences of disease-driving variants.

**Conclusion:** Using MPRA and CRISPRi, JIA risk haplotypes can be queried to identify plausible candidates for disease-driving variants. Once these candidate variants are identified, target genes can be identified using CRISPRi informed by the 3D chromatin structures that encompass the risk haplotypes.

## Introduction

Juvenile idiopathic arthritis (JIA) is a term used to describe a group of childhood illnesses characterized by chronic inflammation and hypertrophy of synovial membranes. Although more rare than the adult disease it resembles, i.e., rheumatoid arthritis (RA), JIA is one of the most common chronic disease conditions in children(1, 2). Like RA, JIA has long been recognized as a complex genetic trait, in which multiple genetic loci contribute to disease risk(3). Although the contribution of any single genetic locus is small, genetic influences on JIA are still quite strong. For example, in a study using the Utah Population Database(4), Sampath et al found that the relative risk for JIA in siblings was nearly 12-fold that of the broader population (11.6; confidence intervals 4.9–27.5; p<3×10^−8^), and that for first cousins was nearly 6-fold (5.8; confidence intervals 2.5–13.8; p<6×10^−5^).

To date more than 30 different risk loci have been shown to contribute to JIA (5-7); the statistical associations are particularly strong for those published in the Hinks Immunochip study(6) and the more recent McIntosh meta-analysis(7). However, the field still faces several challenges in elucidating the mechanism(s) through which genetic variants confer risk, the most formidable of which is to identify the variants that exert the relevant biological effects that contribute to risk. The standard approach – fine mapping, re-sequencing, imputation, bioinformatic annotation, and laboratory testing – is inherently laborious and low throughput due to linkage disequilibrium (LD), which renders causal variants statistically indistinguishable from neutral variants on risk haplotypes. Not knowing the causal variants complicates the task of identifying the target genes. Until the actual causal variants are known, it will be impossible to clarify the mechanisms through which genetic variants alter normal function and contribute to disease risk.

It is becoming increasingly clear that genetic risk for multiple complex traits is more likely exerted through alterations in genomic regulatory functions that influence the efficiency of transcription rather than through the alterations in the coding sequences of disease-relevant genes(8, 9). We have published several papers that support this concept for JIA (10-12). For example, we have shown that the JIA risk loci are highly enriched, compared to genome background, for H3K27me1 and H3K27ac ChIPseq peaks, epigenetic features typically associated with enhancer function(10). Furthermore, we have shown that, *in vitro*, risk variants on the JIA risk haplotypes alter the efficiency of enhancers in the *IL2RA* and *IL6R* risk loci(12). However, the locus-by-locus approach that we and others have used is time-consuming and inefficient. What the field requires is a rapid method for screening thousands of variants for their effects on gene expression in a single assay (13). Here, we report preliminary studies using a massively parallel reporter assay (MPRA) to query SNPs within JIA associated genetic risk loci. We demonstrate the efficacy of this unbiased approach and its capacity for uncovering previously unsuspected mechanisms of genetic risk. Furthermore, we demonstrate how MPRA information and three dimensional chromatin data can then be used to identify the likely target genes, i.e., the genes regulated by the enhancers harboring the variants identified on MPRA. This systematic approach provides an efficient and effective method for resolving the most vexing aspects of autoimmune disease genetics.

## Methods

The general workflow for the MPRA is shown in **Figure 1**.

**Figure 1.**
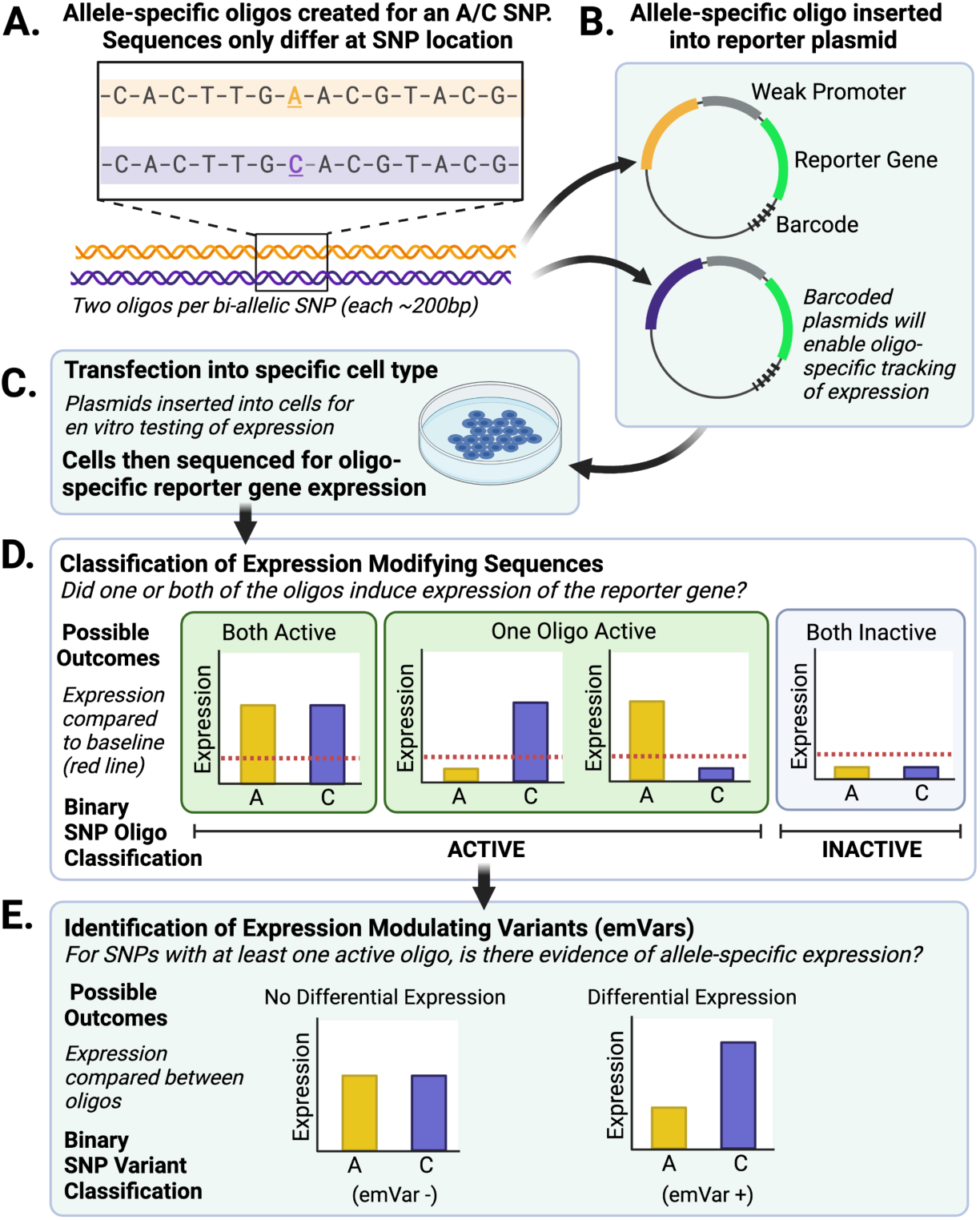
Summary of MPRA work flow and variant screening process. Figure created with BioRender.

We provide further detail in the following sections.

### Selection of genetic variants

We queried SNPs within LD blocks where there is already-established risk for JIA. We used the SNPs identified by Hinks and Hersh (5, 6), as well as 1,016 new SNPs that we identified within the JIA-associated LD blocks using deep whole genome sequencing(14). We used a cut-off of r^2^ = 0.80 to choose those SNPs in strong LD with the index SNPs. This procedure identified 7,312 candidates to test.

### Oligonucleotide library preparation

The oligonucleotide library was prepared following previously published methods(13). In brief, oligos were synthesized (Agilent Technologies) as 230 bp sequences containing 200 bp of genomics sequence and 15 bp of adapter sequence at either end (5’ACTGGCCGCTTGACG [200 bp oligo] CACTGCGGCTCCTGC3’). Unique 20 bp barcodes were added by PCR along with additional constant sequence for subsequent incorporation into a backbone vector by gibson assembly. The oligo library was expanded by electroporation into E.coli and the resulting plasmid library was sequenced by Illumina 2×150 bp chemistry to acquire barcode/oligo pairings. The library underwent restriction digest and GFP with a minimal TATA promoter was inserted by gibson assembly resulting in the 230 bp oligo sequence positioned directly upstream of the promoter and the 20 bp barcode falling in the 3′ UTR of GFP. After expansion within E.coli the final massively parallel reporter assay (MPRA) plasmid library was sequenced by Illumina 1×30 bp chemistry to acquire a baseline representation of each oligo within the library.

### Transfection of K562 cells

Previous work from our group has shown that JIA risk loci are highly enriched for both H3K4me1 and H3K27ac peaks in both neutrophils and CD4+ T cells. Furthermore, for many of the regions queried, H3K4me1/H3k27ac-marked regions were identical in both cell types(10, 11), suggesting that these enhancers regulate common hematopoietic cell functions. This idea is corroborated by ontology analyses of genes within the topologically associated domains (TADs) that encompass the JIA risk haplotypes(12). Thus, we elected to perform our first-step screening using K562 cells, a myeloid cell line that is easy to transfect and has been used previously with this assay(13).

Cells were grown in RPMI medium (Life Technologies) supplemented with 10% FBS (Life Technologies) maintaining a cell density of 0.5-8×10^5^ cells per mL at 37°C and 5% CO2. Libraries were electroporated into K562 cells in 100ul volumes using the Amaxa system (Program X-001, Neucleofactor kit V, Lonza). We performed 6 independent replicates with each replicate consisting of ∼2×10^8^ cells. In a separate set of experiments, K562 cells were treated with or without IFNγ (250 ng/ml) 24 hours after transfection. The cells were collected 48 hours post transfection by centrifugation and washed three times with PBS. The cell pellets were stored at -80°C.

### Preparation of GFP RNA and RNAseq

Total RNA was extracted from cells using RNeasy Midi kit (Qiagen) following the manufacturer’s protocol, including the on-column DNase digestion. A second DNase treatment was performed on the purified RNA using 5 μL of Turbo DNase (Life Technologies) in 300 μL of total volume for 1 hour at 37°C. The digestion was stopped with the addition of 3 μL 10% SDS and 30 μL of 0.5M EDTA followed by a 5 minute incubation at 70°C. The total reaction was then used for pulldown of GFP mRNA. The DNase digested RNA (300 ul) with 300 μL of 20X SSC (LifeTechnologies), 600 μL of Formamide (Life Technologies) and 10 μL of 10 uM biotin-labeled GFP probes (CCTCGATGTTGTGGCGGGTCTTGAAGTTCACCTTG/3BioTEG; CCAGGATGTTGCCGTCCTCCTTGAAGTCGATGCCC/3BioTEG; CGCCGTAGGTGAAGGTGGTCACGAGGGTGGGCCAG/3BioTEG) was incubated for 2.5 hours at 65°C. Biotin probes were captured using 125 μL of pre-washed Streptavidin beads (RNAse clean C1 beads, Life Technologies). The hybridized RNA/probe bead mixture was agitated on a rotator at room temperature for 20 minutes. Beads were captured by magnet and washed once with 1x SSC and twice with 0.1x SSC. Elution of RNA was performed by the addition of 25 μL water and heating of the water/bead mixture for 2 minutes at 70°C followed by immediate collection of eluent on a magnet. A second elution was performed by incubating the beads with an additional 25 μL of water 2 minutes at 80°C. A final DNase treatment was performed in 50 μL total volume using 1 μL of Turbo DNase incubated for 4 hours at 37°C followed by inactivation with 1 μL of 10% SDS and purification using RNA Ampure XP beads (Beckman Coulter).

First-strand cDNA was synthesized from half of the DNase-treated GFP mRNA with SuperScript III and a primer specific to the 3’ UTR (CCGACTAGCTTGGCCGC) using the manufacturer’s recommended protocol.

To minimize amplification bias during the creation of cDNA tag sequencing libraries, samples were amplified by qPCR to estimate relative concentrations of GFP cDNA using 2 μL of sample in a 20 μL PCR reaction containing 10 μL Q5 NEBNext master mix, 2 μL SYBR green I diluted 1:10,000 (Life Technologies) and 0.5 uM of TruSeq_Universal_Adapter (AATGATACGGCGACCACCGAGATCTACACTCTTTCCCTACACGACGCTCTTCCGATCT) and MPRA_Illumina_GFP_F primers (GTGACTGGAGTTCAGACGTGTGCTCTTCCGATCTCGCCCTGAGCAAAGACC). Samples were amplified with the following conditions: 95°C for 20 seconds, 40 cycles (95°C for 20 sec, 65°C for 20 sec, 72°C for 30 sec), 72°C for 2 min.

To add Illumina sequencing adapters, cDNA samples and 4 mpra:gfp plasmid controls were diluted to match the replicate with the lowest concentration and 20 μL of normalized sample was amplified using the reaction conditions from the qPCR scaled to 50 ul. Amplified cDNA was purified using MinElute PCR Kit (Qiagen) and eluted in 30 μL of EB. Individual sequencing barcodes were added to each sample by amplifying the 20 μL elution in a 50 μL Q5 NEBNext reaction with 0.5 uM of TruSeq_Universal_Adapter primer and a reverse primer containing a unique 8 bp index (Illumina_Multiplex) for sample demultiplexing post-sequencing. Samples were amplified at 95°C for 20 seconds, 6 cycles (95°C for 20 sec, 64°C for 30 sec, 72°C for 30 sec), 72°C for 2 minutes. Indexed libraries were purified using MinElute PCR Kit (Qiagen) and pooled according to molar estimates from Agilent TapeStation quantifications. Samples were sequenced using 1×30 bp chemistry on an Illumina HiSeq through the Jackson laboratory in Maine.

### Data analysis: Identification of SNPs with significant influences on gene expression

We followed the same analysis used previously(13) on our MPRA data from K562 cells. For each of the replicates, we extracted the first 20bps from the sequenced reads and used them to assign the reads to either reference or alternative allele for variants based on our barcode library. We then generated a count-table for each variant, including the names of variants, as well as the counts of the reference and alternative alleles for each replicate. After merging replicates, we generated a master count-table for each of the variant alleles for each of the replicate of DNA (plasmid library), RNA expression in K562 cells, and RNA expression in K562 cells with IFNγ treatment. We used a customized R script based on that described by Tewhey(13) to process the count-table and identify the DNA elements, defined as a region on chromosome that is investigated in MPRA, with active regulatory activities and the variants that can alter the regulatory activities. We used DESeq2(15), to normalize the counts and fit the dispersion using ‘local fit’. The distribution of the log2 fold changes between RNA and DNA were investigated. Since we found the distribution was not centered at 0, we adjusted the size factors for DNA and RNA samples according to the offset, then performed the normalization again. The oligos showing differential RNA expression relative to the plasmid DNA were then identified by the nbinomWaldTest function in DESeq2 (a Wald Test for the coefficients in a negative binomial generalized linear model) and applying a threshold of 0.01 for the False Discovery Rate and a minimum fold change of 1.5. For DNA elements that displayed significant regulatory activity, we applied a t-test on the log2-transformed RNA/plasmid ratios for each paired replicate (e.g. alternative alleles of K562 replicate 1 vs reference alleles of K562 replicate 1) to test whether the reference and alternate allele had a similar activity. The p-values from t-test were adjusted through the Benjamini-Hochberg process into FDR values and we applied a cutoff of FDR 0.01 to call those variants showing altered regulatory activities between alleles.

### Development of K562 cells that express dCas-KRAB

The inducible dCas9-KRAB construct was generated as described previously(16-18). The dCas9-KRAB were co-transfected with SB100X into K562. The dCas9-KRAB was integrated into genome of K562 cells by SB100X, a hypersensitive transposon(19). Cells were selected using blasticidin for 10 days. These cells express dCas9-KRAB in a doxycycline-dependent manner (**Supplementary Figure 1**).

**Supplementary Figure 1.**
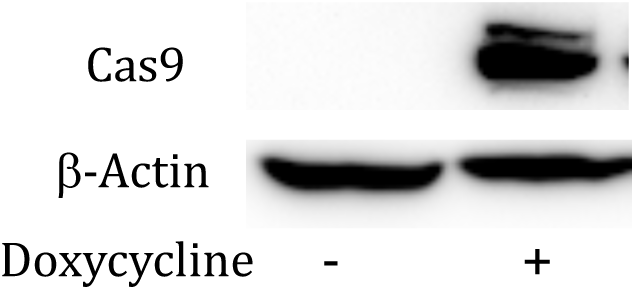
Western blot shows that the Cas9 protein induced by doxycycline in K562 cells transfected dCas9-KRAB construct after treated with 1 ug/ml of doxycycline for 48h.

### Identification of genes regulated by enhancers harboring SNPs that screen positive on MPRA

Because most enhancers regulate genes within the same chromatin loop or *topologically associated domain* (TAD)(20), we used publicly available HiC data(12) to identify the likely target genes regulated by the enhancers harboring the SNPs identified by MPRA. As proof-of-concept, we chose to test genes from 2 loci. We chose *TRAF1*, which is of special interest because of its specific association with a polyarticular disease course(21), and *ERAP2/LNPEP*, which are downstream effectors of interferon responses(22), which are known to be enhanced in JIA(23).

Single guide RNAs (sgRNAs) were designed by using a web service from IDT Integrated DNA Technologies, Inc. (www.idtdna.com/site/order/designtool/index/CRISPR_CUSTOM). We designed four gRNAs for each enhancer (**Figure 2**). The sgRNA sequence for the HS2 enhancer (HS Cr4, gaaggttacacagaaccaga) was from (24). The gRNA expression vectors were generated as described previously(16-18). The sequence information of all constructs was verified by Sanger sequencing.

**Figure 2.**
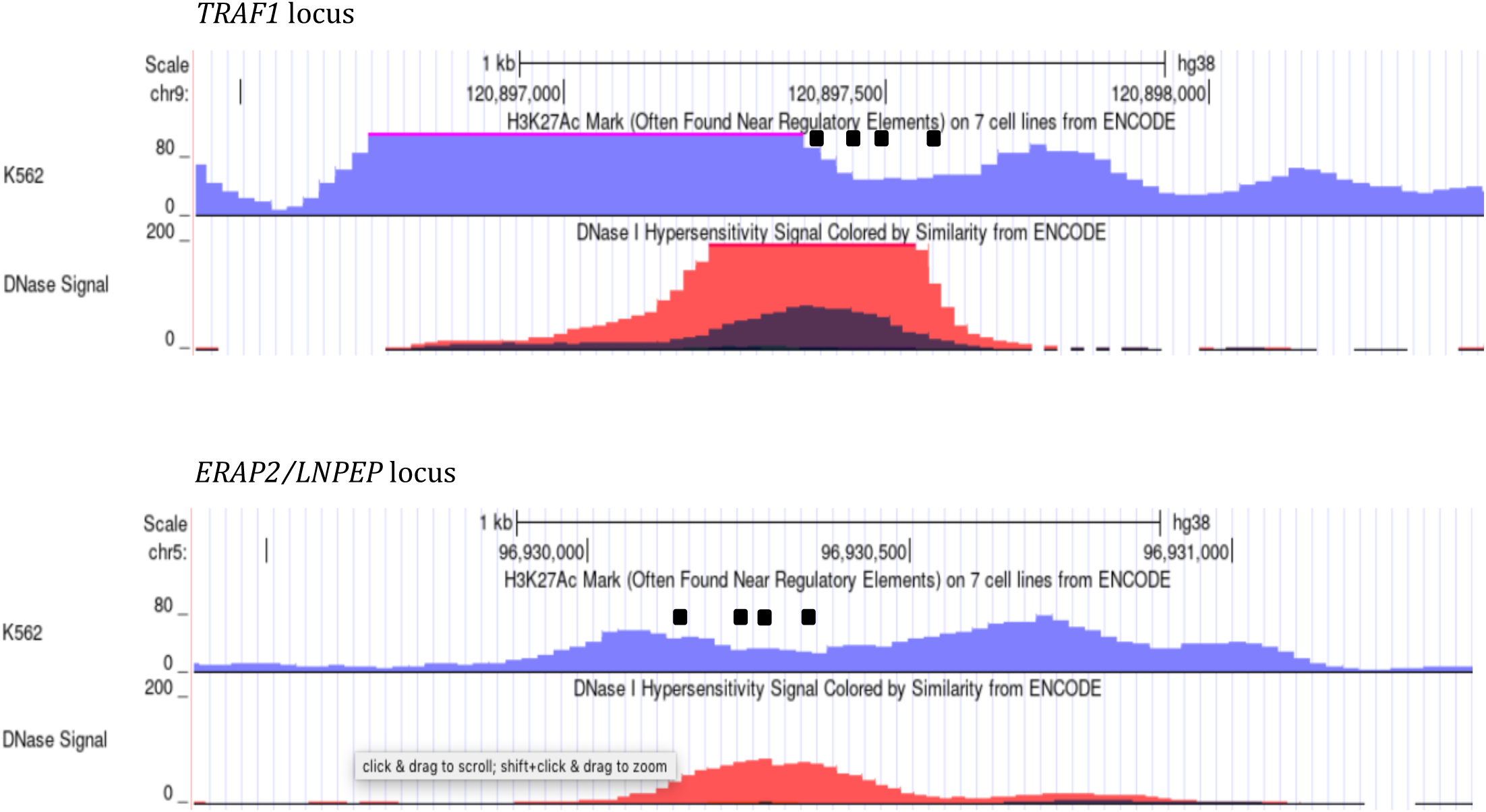
Genome browser screen shots showing the positions of gRNAs used to interrogate enhancer regions using CRISPRi. An intergenic enhancer region in the TRAF1 locus is shown in the upper panel, and an enhancer in intergenic region in ERAP2/LNPEP is shown in the lower panel. gRNAs are shown as black rectangles. The purple peaks are derived from ENCODE H3K27ac ChIPseq data and the red peaks from ENCODE DNaseI hypersensitive data, both in K562 cells.

Next, a pool of gRNAs (4 gRNAs) was transfected into the dCas9-KRAB K562 cells using hypBase plasmid(25), which inserts the gRNA into the genome of the dCas9-KRAB-expressing K562 cells. Cells were selected with puromycin for 10 days to get stably expressed gRNAs-dCas9-KRAB K562 cells. To induce dCas9-KRAB expression, the cells were treated with 1 ug/ml of doxycycline for 48h before RNA isolation.

### RT-qPCR

Total RNA was purified using RNeasy Plus mini kit (Qiagen), and cDNA was synthesized by with iScript™ cDNA synthesis kit according to the directions of the manufacturer (Bio-Rad, Hercules, CA, USA). Quantitative PCR was performed using Power SYBR™ Green PCR Master Mix kit on a StepOne Plus real time PCR system (Applied Biosystems, Foster City, CA, USA). The relative expression level of a gene was normalized by that of GDPDH. The primer sequences were, PHF19: forward- TGGACAGATGGCCTGTACTA and reverse- CCTTCCATAGGACCCAGTATTT; LNPE: forward- TGAGCAATACACCGCTTTATCA, reverse- GTGCTCATCTTCACACTCTCAG; ERAP2: forward- GACCTCTTCTGCTTCCGATAAA, reverse- GCCAAATATCATTCCACCATTCC; CAST: forward- TGACCGGTCTGAATGTAAAGAG, reverse- TATACTACACATGGAGGTCCGA; HBE1: forward- TCACTAGCAAGCTCTCAGGC, reverse- AACAACGAGGAGTCTGCCC.

## Results

### MPRA screening identifies multiple variants with transcriptional effects

We transfected K562 cells with oligonucleotide probes representing 7,312 SNPs on the JIA risk haplotypes identified from GWAS and candidate gene studies (5, 26) as well as the Immunochip(6). After final quality control measures, there were 5,226 sequences with sufficient representation in the MPRA library to undertake downstream analyses. Of these, 1,482 (28%) showed regulatory activity. Of these, 530 (19.8%) showed a significant difference from the common allele in unstimulated K562 cells, and 490 (18%) showed differential expression in IFNg-stimulated cells, using FC > 2 and FDR =0.01 as a cut-off. After excluding SNPs within the HLA class I and class II loci (n=406), where coding functions are believed to be the most important disease drivers, we further filtered SNPs to identify those that were in open chromatin and within H3K27ac ChIPseq peaks (unstimulated cells) or open chromatin and H3me1, but not H3K27ac-marked regions for SNPs identified in exclusively in stimulated cells. Using these procedures, we identified n=42 SNPs in unstimulated K562 cells (**Table 1)**. After stimulation with IFNγ (250 ng/ml), we identified an additional 42 SNPs that showed significant effects on gene expression that were not identified in unstimulated cells (**Table 2**). These findings are consistent with previously published studies that suggest that many disease-relevant alleles may exert their effects on immune cells only after those cells are activated(28).

**Table 1.**
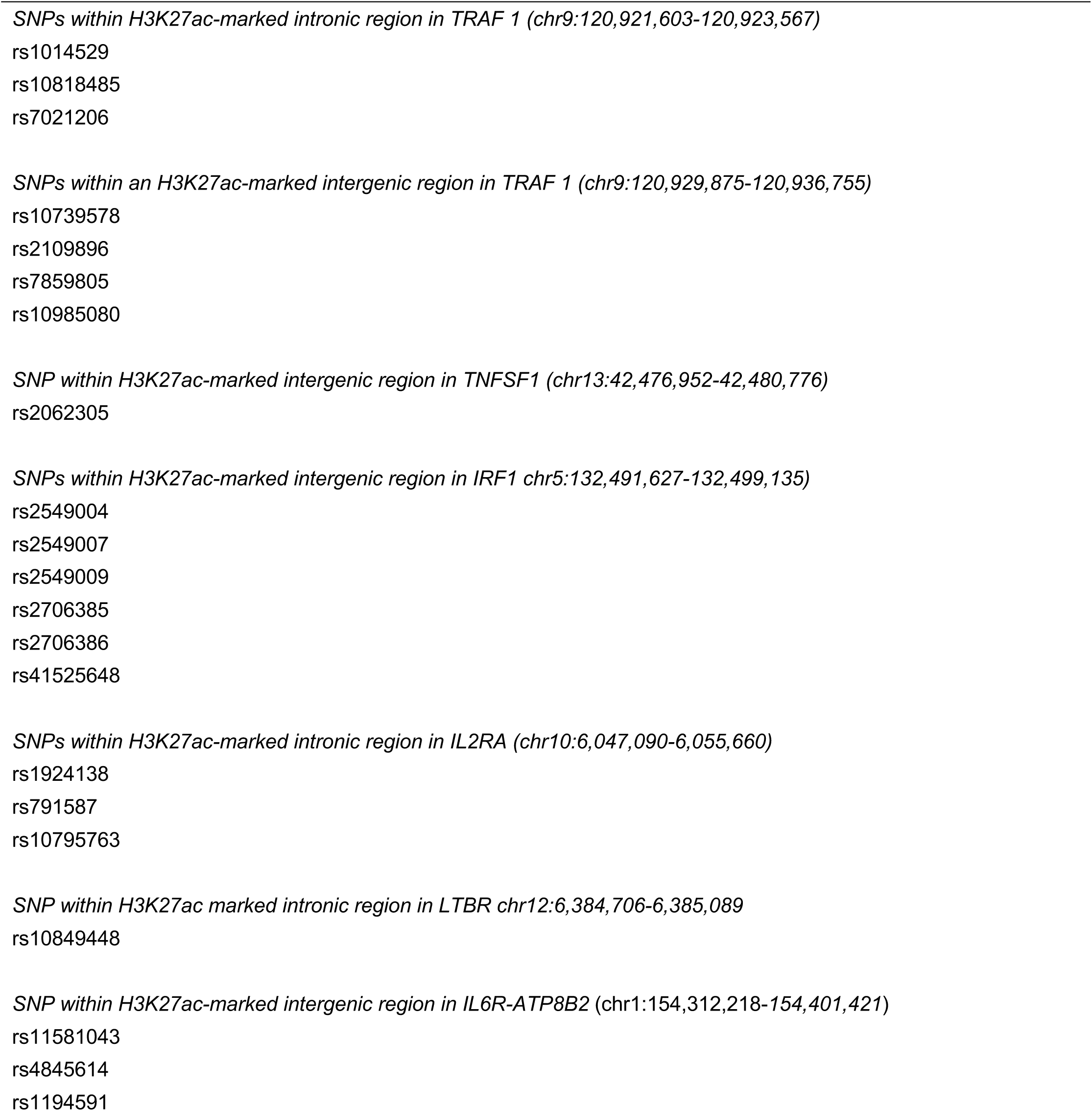
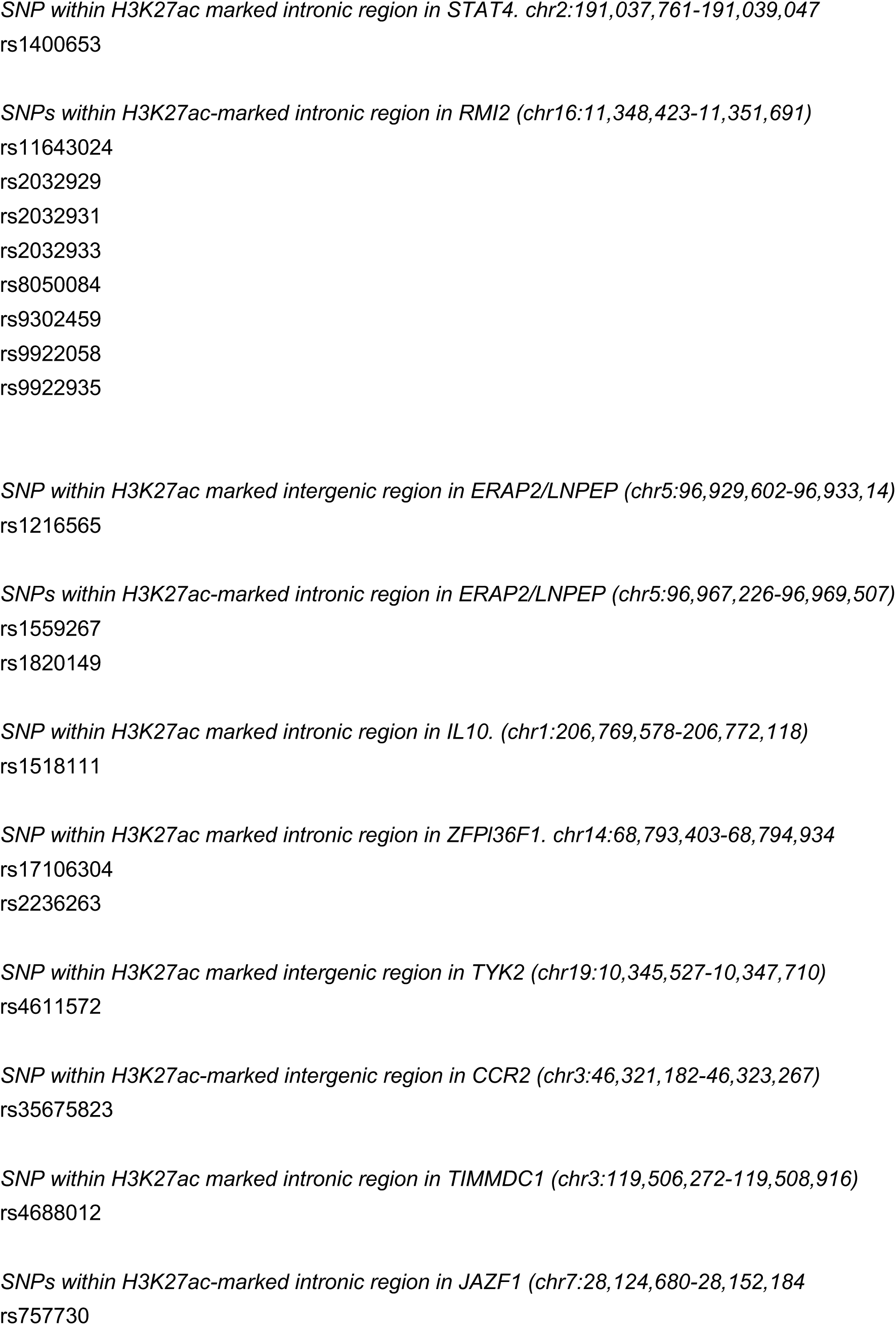
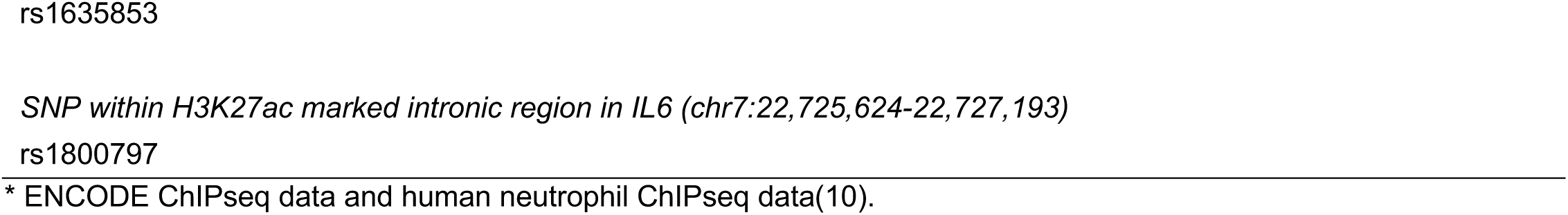
SNPs Screening Positive in Unstimulated K562 Cells (n=42)

**Table 2.**
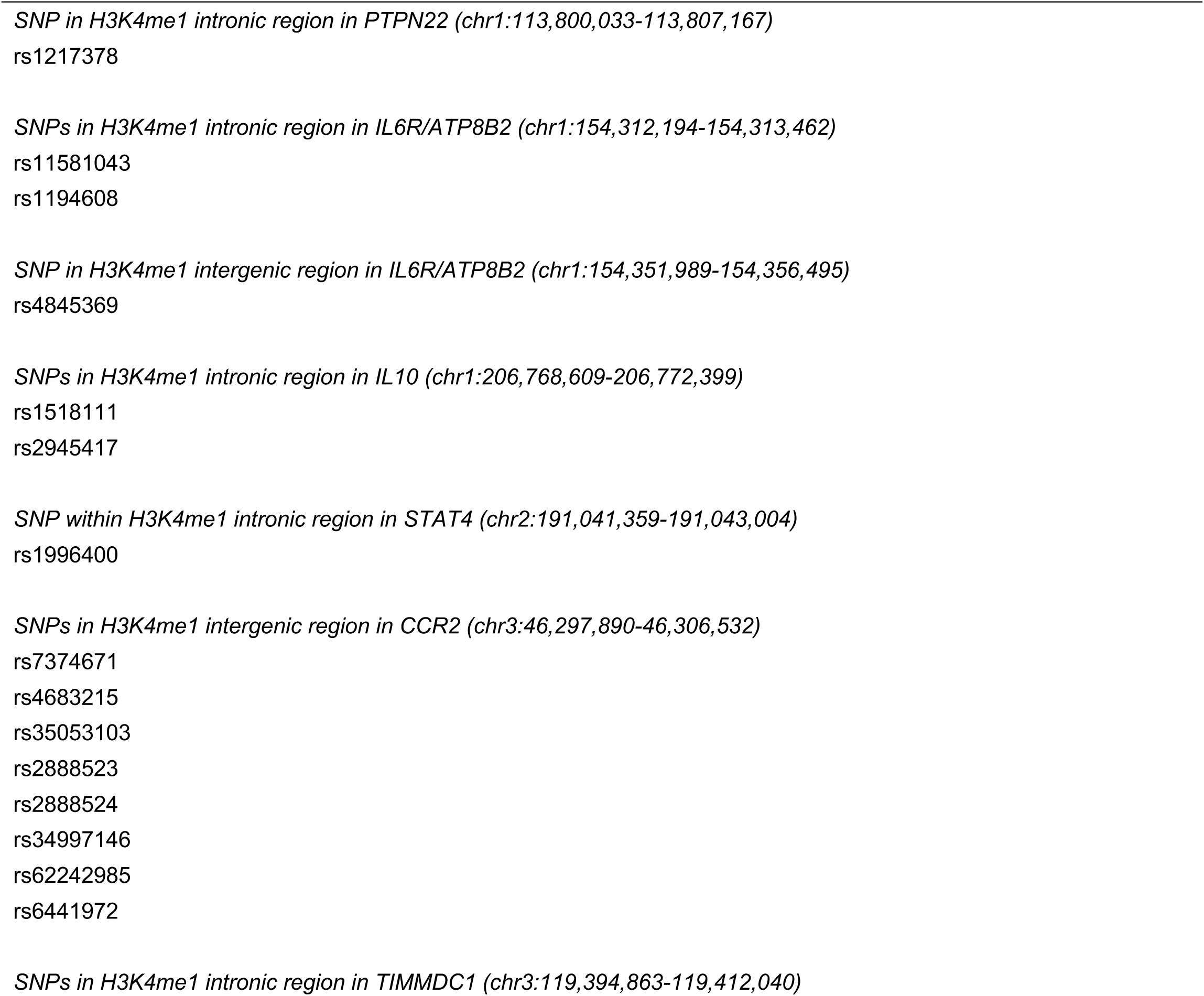
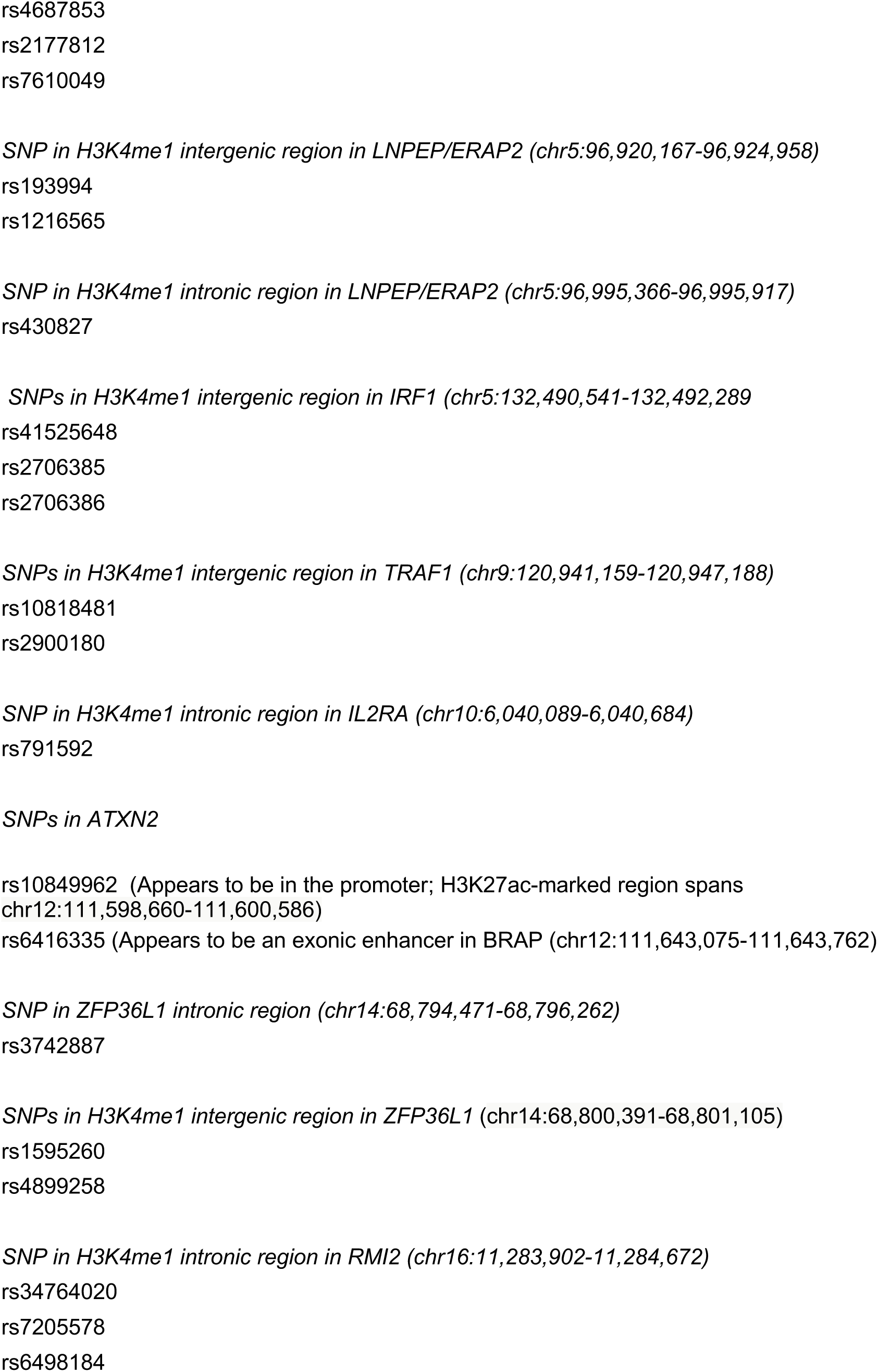
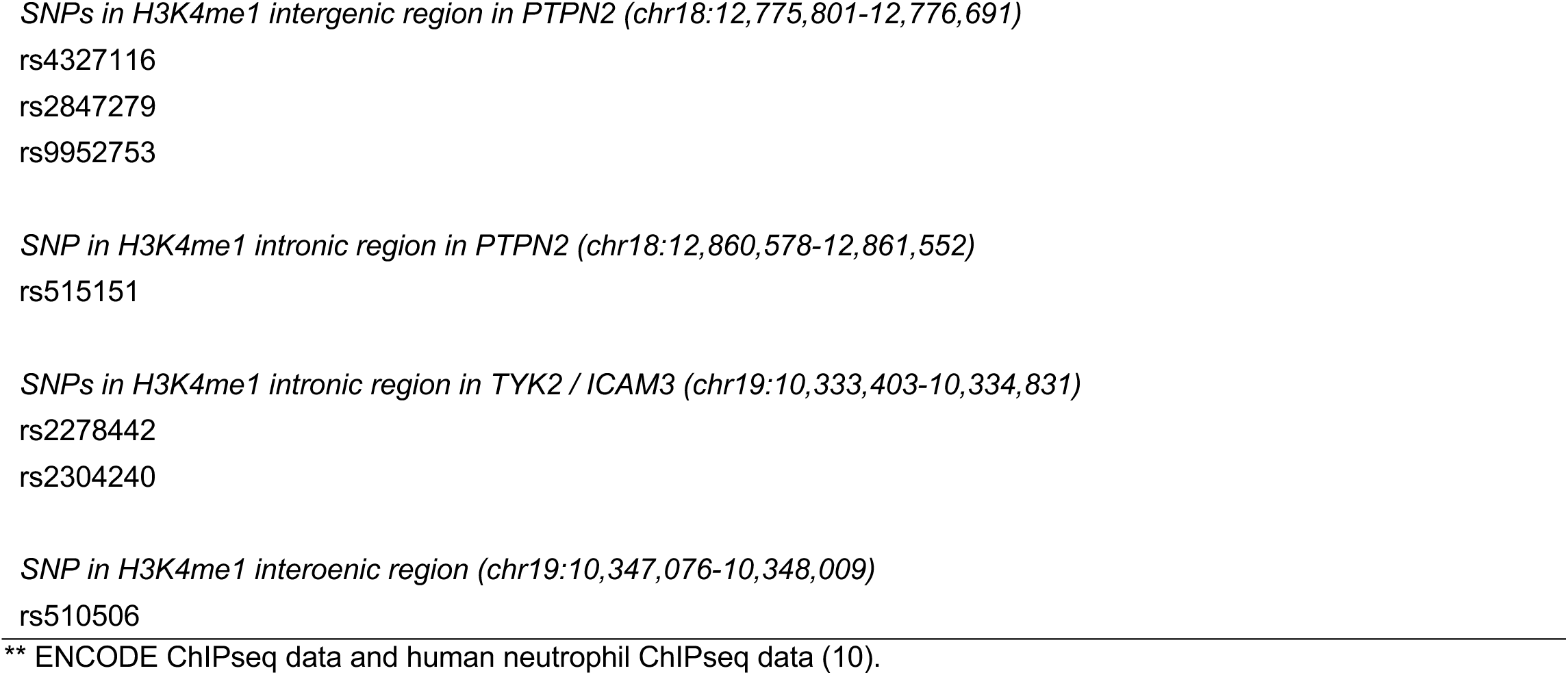
SNPs Screening Positive in K562 Stimulated By Interferon Gamma (n=42)

In many cases, we identified multiple alleles on the same haplotype, although these alleles were not necessarily within the same functional element. For example, in the *TRAF1, LNPEP/LNPEP/ERAP2*, and *IL6R/ATP8B2* loci, we identified variants within both intergenic and intronic enhancers in unstimulated K562 cells. In IFNγ-stimulated cells, we identified variants in both intergenic and intronic enhancers in the *IL6R/ATP8B2, TYK2/ICAM3*, and *LNPEP/ERAP2* loci. In stimulated K562 cells, we also identified expression-enhancing variants that are situated within the promoter and within an exon within the *ATXN2* locus. This finding may reflect that these regions have enhancer as well as promoter/coding functions, as has been described for other genomic regions(29, 30).

### Identification of target genes

The identification of risk-enhancing alleles of JIA haplotypes may facilitate the identification of target genes, i.e., the genes whose expression levels are influenced by the risk-driving SNPs. To test this concept, we relied on the fact that most enhancers regulate genes within the same TAD(20) and that the 3D chromatin structures are identifiable from publicly available chromatin conformation data.

We tested the intergenic enhancer harboring rs10985080, located at chr9:120,896,106-120,897,428 (GRCh38/hg38) on the *TRAF1* haplotype, as well as the intergenic enhancer on the *LNPEP/ERAP2* harboring rs1216565 as described in the *Methods* section. Using publicly available HiC data from the *3D Genome Browser* (http://3dgenome.fsm.northwestern.edu(31)), we visualized heat maps defining TAD structures in K562 cells and reported by Rao et al(32). We identified genes within the same chromatin loops that were likely targets of these enhancers. We show HiC data derived from K562 cells in **Figure 3**. For *TRAF1*, we identified *TRAF1, PHF19* and *C5* as the most likely candidate targets for this enhancer.

**Figure 3.**
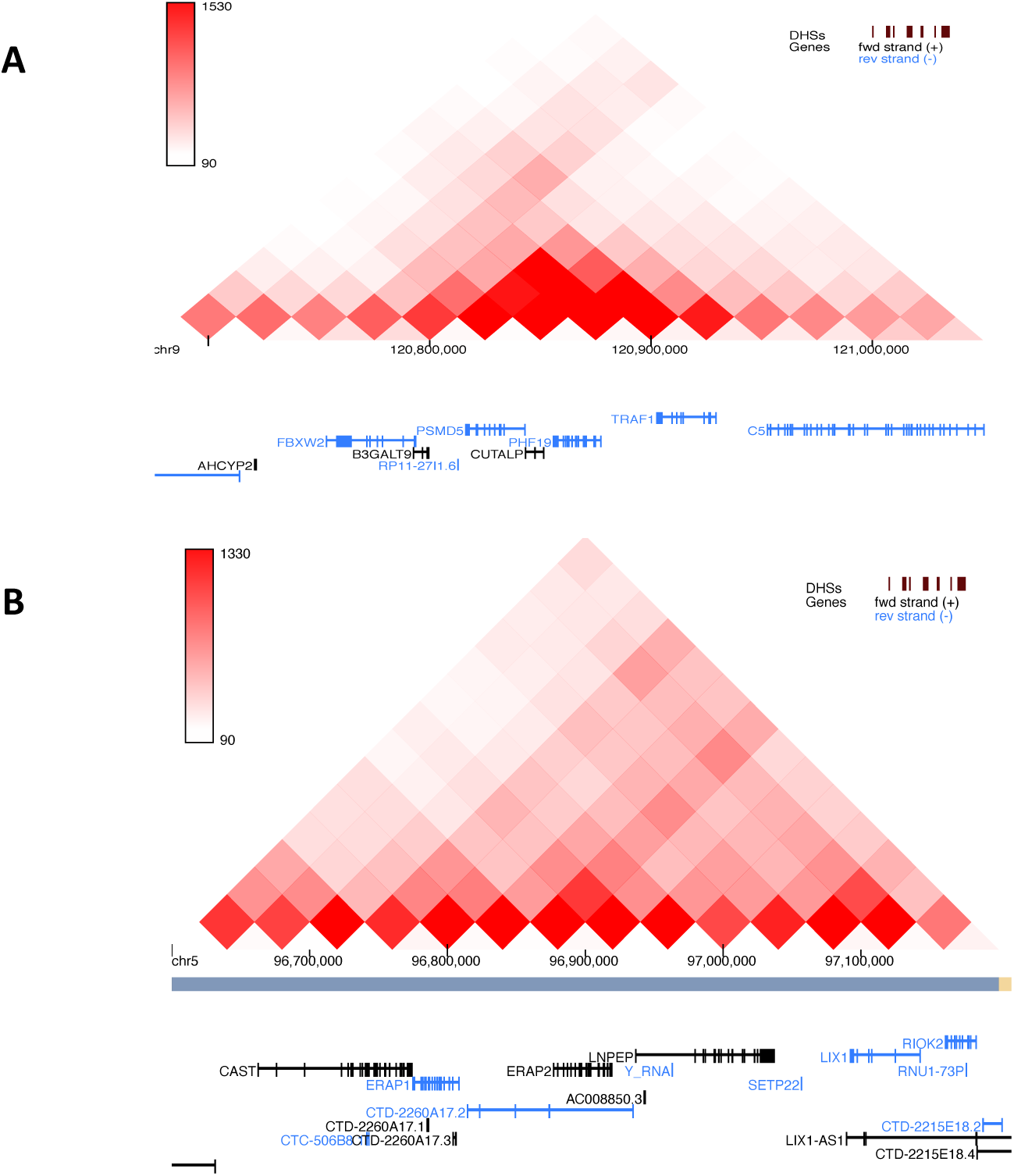
Heat map derived from HiC analysis of K562 cells showing the 3D chromatin structure around the TRAF1 locus (A) and LNPEP/ERAP2 (B). Genes in black are those on the forward strand, while those in blue are on the reverse strand. Data are derived from Rao et (32). Note that not all the genes in this TAD are expressed in K562 cells. For example, we detected 3 genes (PHF19, TRAF1 and C5), which located in TRAF1 locus, by qPCR, only PHF19 and C5 were expressed in K562 cells. Note that TRAF1 itself was not.

Within the TAD encompassing the intergenic enhancer in the *TRAF1* locus we identified 2 expressed genes, *PHF1* and *C5*. Note that *TRAF1* itself was not expressed. Attenuation of the *TRAF1*intergenic enhancer by CRISPRi resulted in a significant (p<0.01) reduction in expression of the *PHF19* gene, but not *C5* (data not shown). These results are shown in **Figure 4**. We next examined the intergenic enhancer harboring rs1216565, located at chr5:96,929,854-96,931,091 (GRCh38/hg38) in the *LNPEP/ERAP2* locus.

**Figure 4.**
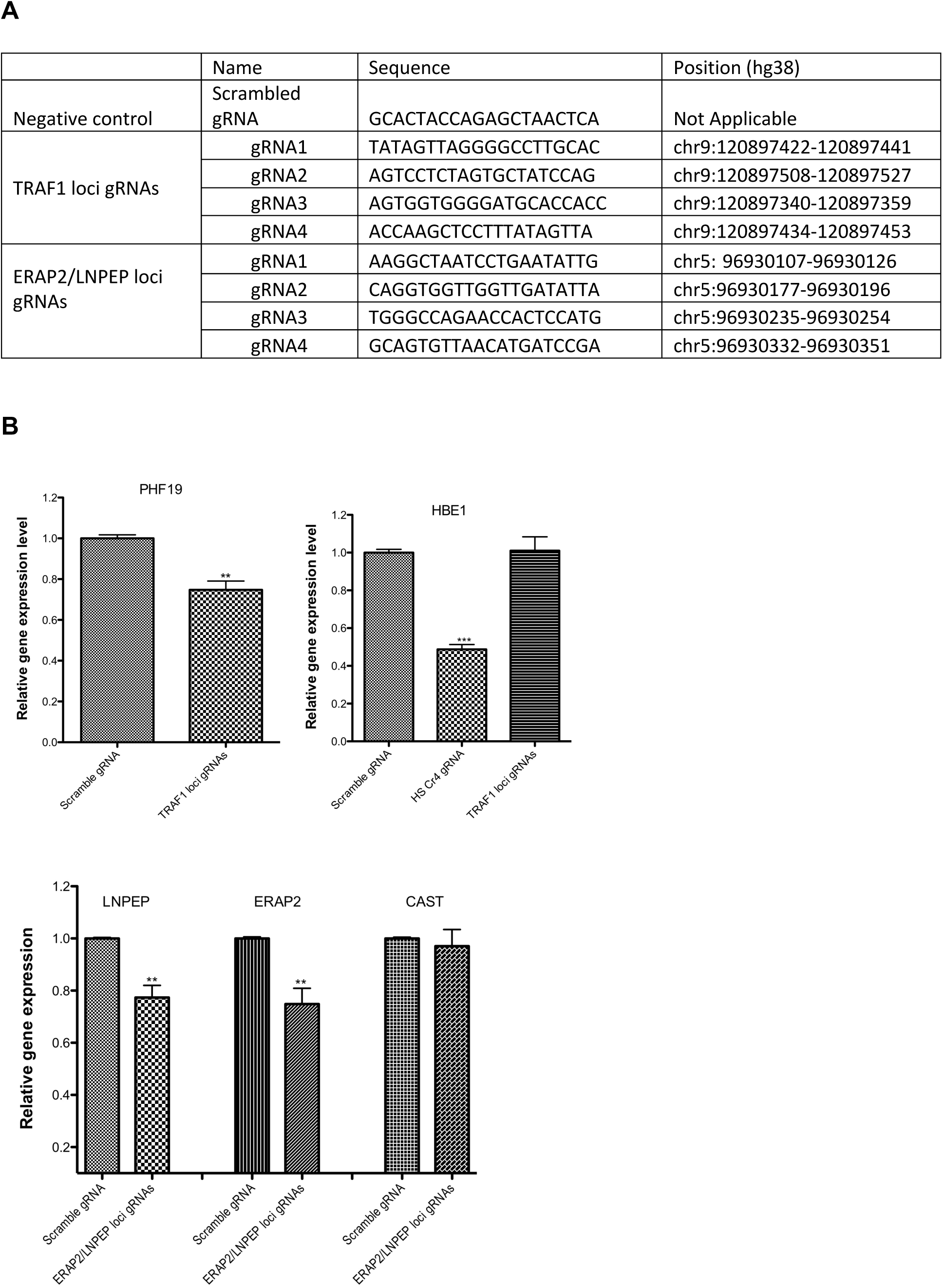
CRISPi identifies target genes of enhancers harboring JIA-associated variants detected on MPRA. (**A**). gRNAs used for epigenome editing experiments. The scrambled gRNA is a non-targeting negative control gRNA. For each experiment, 4 gRNAs were targeted to the functional regions of intergenic enhancers in the TRAF1 locus and the ERAP2/LNPEP locus. **(B)** Attenuation of the intergenic enhancer at the TRAF1 locus significantly reduced expression of PHF19, but not C5 (not shown). Off-target effects are monitored by showing that these gRNAs have no effect on HEB1 expression, but that gRNAs directed to an enhancer known to regulated HEB1 attenuates expression. Bar graphs summarize the results of 4 independent experiments. **(C)** Attenuating the intergenic enhancer in the LNPEP/ERAP2 locus reduces expression of both LNPEP and ERAP2, but not the adjacent gene, CAST, nor the HEB1 gene (not shown). In each experiment, scrambled versions of the gRNAs had no effect on expression of the putative targets. **p<0.01; ***p<0.001

Once again, we used publicly available 3D chromatin data to identify likely targets, which included *LNPEP, ERAP2*, and *CAST*. As shown in **Figure 4C**, attenuation of this enhancer resulted in significant reductions in the expression of both *LNPEP* and *ERAP2*, but not the adjacent gene, *CAST*. This finding is consistent with the known capacity of enhancers to regulate multiple genes within the same topologically-associated domain (20).

Finally, we sought to gain additional information that MPRA-identified SNPs located in the enhancers in the *TRAF1* and *LNPEP/ERAP2* locus influence gene expression of the candidate target genes in humans. We used the Genotype-Tissue Expression (GTEx) project’s (33) expression quantitative trait locus (eQTL) calculator and querying whole blood expression data for this purpose (https://www.gtexportal.org/home/testyourown). Results of these analysis are shown in **Table 3**, below. SNPs highlighted in bold indicate those with greater-than-by chance likelihood that the SNP has an influence on the expression of the listed gene.

**Table 3.** MPRA-identified SNPs located in the enhancers in the *TRAF1* and *LNPEP/ERAP2* loci influence gene expression of the candidate target genes in humans.

| Locus | Gene | SNP | P-value | P-value threshold | NES* |
| --- | --- | --- | --- | --- | --- |
| *TRAF1* | *PHF19* | rs10739578 | 0.00077 | 0.00021 | 0.065 |
|  |  | rs2109896 | 0.084 | 0.00021 | 0.036 |
|  |  | rs7859805 | 0.035 | 0.00021 | 0.044 |
|  |  | **rs10985080** | **0.000068** | 0.00021 | 0.078 |
| *LNPEP/ERAP2* | *ERAP2* | **rs1216565** | **2.4e-152** | 0.00020 | 0.87 |
|  | *LNPEP* | **rs1216565** | **0.0000064** | 0.00020 | 0.072 |

NES = Normalized effect size

In each case, then, GTEx whole blood expression data further strengthens the predictions from the MPRA + CRISPRi analysis. At the same time, these analyses demonstrate the utility of our approach in clarifying GTEx data where multiple SNPs in strong LD appear to influence the expression of a gene. Furthermore, when multiple expression-altering alleles are adjacent (and therefore in strong LD), the GTEx analysis may clarify the specific SNP(s) that exert the strongest influence on the candidate target genes. Note that for rs10985080 and rs1216565, despite significant differences from the common allele, the effect sizes are relatively small. This finding is consistent with the observations of Gasperini et al (20), who have shown that the effect sizes for most enhancers on the genes they regulate is in the range of 15-30%.

## Discussion

The field of genetics as applied to complex traits has started to move beyond the identification of genetic associations and toward the elucidation of the mechanisms through which genetic variants confer risk (34, 35). However, a significant impediment to accomplishing this task is the fact that a strength GWAS studies, which leverage LD to identify regions conferring genetic risk, is also a weakness, in that the SNPs that tag genetic risk loci are in LD with dozens, sometimes hundreds, of other SNPs, most of which have no influence at all on disease risk. Thus, distinguishing the true causal variants (i.e., those that exert the relevant biological effects) from the innocuous ones in which they are in LD, has been a challenge. At the same time, the discovery that, for most complex traits (8), including autoimmune diseases (9), genetic risk is likely to impinge on regulatory functions rather than the protein-coding sequences of pathology-driving genes, has complicated the search for target genes (i.e., the genes influenced by the causal variants).

In this paper, we demonstrate a systematic strategy for identifying both causal variants and their target genes on JIA risk haplotypes. We find that, by using existing chromatin data in combination with MPRA to screen for expression-altering variants, we can identify a finite number of variants that, based on their functional properties, are strong candidates as actual causal variants, as others have recently shown(27). Subsequent identification of target genes can then be accomplished using CRISPRi approaches, especially for those variants that lie within enhancers, which are likely to be fundamental to autoimmune disease pathogenesis(36). The CRISPRi experiments are simplified by the fact that most enhancers regulate genes within the same TAD(20), and, thus experiments can be performed in a targeted fashion rather than genome-wide. Finally, one can make a causal link between expression levels of genes identified by CRISPRi and the variants that screen positive on MPRA using GTEx whole blood expression data.

The MPRA screening yielded some surprising results. We note, for example, that there were many loci where we identified multiple expression-altering alleles within the same functional element. For example, rs2549004, rs2549007, rs2549009, rs2706385, rs2706386 and rs41525648 within a single intergenic enhancer on the *IRF1* haplotype and rs1559267, rs1820149, and rs1559267 within a single intronic enhancer on the *TRAF1* haplotype. This finding suggests that the disease-associated haplotypes exert risk because they contain multiple alleles in strong LD that, together, alter immune regulatory functions. Furthermore, many of the risk haplotypes contain more than one functional element that is affected, and these different functional elements may exert their effects on different genes. There is no reason to assume, for example, that the intergenic and intronic enhancers on the *TRAF1* haplotype regulate the same genes.

Another useful fact that emerges from our data is that physical interactions between enhancers and promoters do not necessarily indicate a regulatory relationship. For example, HiC and promoter capture HiC data demonstrate physical interactions between the intergenic enhancer in *LNPEP/ERAP2* and the promoter *CAST* gene. However, attenuating this enhancer had no effect on *CAST* expression. While this finding doesn’t exclude the possibility that this enhancer might work in concert with others to regulate *CAST*, it serves as a precautionary message in how we use and interpret 3D chromatin data and gene expression data from patient cells.

Our work also highlights the utility of using the *Sleeping Beauty* transposase system and hypBase vectors for gRNAs in functional genomics experiments. Genome-wide CRISPRi screens for enhancer activity have typically use lentivirus and/or plasmid vectors to attenuate enhancer function(20). However, these assays can be vexing to perform and replicate because of the low multiplicity of infection (MOI) rates seen with such vectors. This makes it difficult to interpret different experiments or even compare replicates within a single experiment. Our approach allows stable and high levels of expression of both the epigenome editing enzyme and the gRNAs, facilitating both replication and inter-experimental comparisons (e.g., between the efficiency of intronic vs intergenic enhancers in regulating a specific gene).

This work has several limitations, the most important of which is the use of cell lines rather than primary human cells for these assays. There is accumulating data that, based on the similarity of their chromatin to the cognate primary human cells such as Jurkat and THP-1 are suitable as models in genetic studies of human autoimmune diseases(37). However K562, which were derived from a human myelogenous leukemia, are less like their primary human counterparts, although the TAD structures strongly resemble primary myeloid cells(12). Recent developments, which include using a Rous sarcoma virus promoter instead of the minimal promoter we used here, suggest that MPRA assays can be performed in primary human cells, provided that a sufficient number of replicates are performed to attenuate the differences between individual donors(38). We now have such experiments under way in our laboratory.

Another limitation of the findings here was the agnostic nature of the variant selection process. We chose variants with MAF >1% on the JIA haplotypes identified in several different studies (6), following the approach of Tewhey et al(13). Variants were not selected based on their presence within plausible functional chromatin or their frequency in genotyped patients with JIA. As Lu and colleagues have shown(27) using such selection methods can increase the efficiency of the MPRA.

Another important limitation to this approach is the fact that MPRA may not detect effects as they would occur in native chromatin (35, 39). This limitation may lead to both false positives and false negatives. False positives may be reduced by using an additional criterion (or criteria) to filter MPRA-identified variants. For example, Ainsworth et al (40) have shown that applying analyses of DNA topology, an important determinant of DNA non-coding functions, can improve the predictive value of variants screened on MPRA, since these analyses detect a feature intrinsic to native chromatin. Thus, the addition of a filtering feature that detects effects in native chromatin is likely to significantly reduce the number of false positives that emerge from MPRA alone. It should be noted that false negatives, which DNA topology analyses won’t solve, is much less of a problem for the field. The JIA risk haplotypes contain >13,000 SNPs in LD with the SNPs used to identify/tag the risk loci. The urgent need is to reduce this number so that further functional characterization can proceed in an efficient way.

Even after this process is completed, there will be further work to be done. The agnostic nature of GWAS and genetic fine mapping studies makes it impossible to identify the cells whose functions are affected by disease-driving variants. In JIA, there are likely to be multiple cells types, in addition to CD4+ T cells and neutrophils(10, 11) that are influenced by genetic variants. These include B cells(41) and monocytes(42), and possibly other cells that regulate the innate immune response, such as hepatocytes(43). All of these cell types will need to be tested separately, and it is likely that different SNPs will be found to alter different regulatory regions in each cell type.

For many of these cell types, there may be serious limitations in using GTEx whole blood expression data to make the causal connection between SNPs identified on MPRA and expression levels of genes identified on CRISPRi. We note that our MPRA and CRISPRi studies were performed in myeloid K562 cells, and that neutrophils (derived from myeloid precursors) are the most abundant leukocyte in adult peripheral blood. Gene expression profiles from whole blood are strongly influenced by neutrophil expression signatures(44, 45). It seems plausible that genetic influences that specifically affect gene expression in lymphocytes and lymphocyte subsets would be difficult to corroborate using existing GTEx data, especially given the relatively small number of genotyped subjects (fewer than 700) currently in the GTEx whole blood data set. In these cases, establishing a causal link between a given SNP and the expression of candidate target gene might require direct interrogation of the cells of interest using CRISPR/homology-directed repair strategies.

In conclusion, we describe a systematic approach for identifying both causal variants and their target genes on JIA risk haplotypes. This approach relies on knowledge of the chromatin structures that encompass the risk haplotypes as well as the massively parallel genomic assays to identify functional features that make them strong candidates for being disease-driving variants. The use of screening assays in primary human cells and the adaptation of informative cell lines can be expected to rapidly advance our understanding of genetic mechanisms that drive JIA risk.

## Acknowledgements

The authors wish to thank Joshua J. Breunig for providing us the hypBase vectors used in the gRNA transfections. We also that Hannah Ainsworth, PhD, for producing *Figure 1*.

## Author contributions

KJ – Performed the experimental work and assisted in data analysis and interpretation and manuscript presentation.

TL – Assisted in the design of the MPRA library and analyzed the MPRA data. Assisted in data analysis and interpretation and manuscript presentation.

SK – Assisted in the preparation and quality control measures on the MPRA library.

RT – Assisted in the design and quality control of the MPRA library, assisted with data analysis and interpretation and manuscript preparation.

DK and YP – Provided technical advice and guidance for the CRISPRi studies. YP – Assisted in manuscript preparation

JNJ – Designed the study and assisted in the analysis and interpretation of the data. Assisted in manuscript preparation.

